# Comparison of mouse models reveals a molecular distinction between psychotic illness in PWS and schizophrenia

**DOI:** 10.1101/2021.01.26.428232

**Authors:** Simona K. Zahova, Trevor Humby, Jennifer R. Davies, Joanne E. Morgan, Anthony R. Isles

## Abstract

Prader-Willi Syndrome (PWS) is a neurodevelopmental disorder caused by mutations affecting paternal chromosome 15q11-q13, and characterized by hypotonia, hyperphagia, impaired cognition and behavioural problems. Psychotic illness is a challenging problem for individuals with PWS and has different rates of prevalence in distinct PWS genotypes. Previously, we demonstrated behavioural and cognitive endophenotypes of relevance to psychiatric illness in a mouse model for one of the associated PWS genotypes, namely PWS-IC, in which deletion of the imprinting centre leads to loss of paternal imprinted gene expression and over-expression of *Ube3a*. Here we examine the broader gene expression changes that are specific to the psychiatric endophenotypes seen in this model. To do this we compared the brain transcriptomic profile of the PWS-IC mouse to the PWS-cr model that carries a deletion of the PWS minimal critical interval spanning the snoRNA *Snord116* and *Ipw*. Firstly, we examined the same behavioural and cognitive endophenotypes of relevance to psychiatric illness in the PWS-cr mice. Unlike the PWS-IC mice, PWS-cr exhibit no differences in locomotor activity, sensory-motor gating, and attention. RNA-seq analysis of neonatal whole brain tissue revealed a greater number of transcriptional changes between PWS-IC and wild-type littermates, than between PWS-cr and wild-type littermates. Moreover, the differentially expressed genes in the PWS-IC brain were enriched for GWAS variants of episodes of psychotic illness but, interestingly, not schizophrenia. These data illustrate the molecular pathways that may underpin psychotic illness in PWS and have implication for potential therapeutic interventions.

## INTRODUCTION

Prader-Willi syndrome (PWS), is a neurodevelopmental disorder primarily caused by mutations affecting paternally expressed genes (PEGs) on the imprinted 15q11.2-q13 region. This interval includes several genes expressed only from the paternal chromosome (*MKRN3, MAGEL2, NECDIN, NPAP1, SNURF-SNRPN*, cluster of non-coding RNAs), as well as two genes expressed only from the maternal chromosome (*UBE3A, ATP10*)^1^. The disruption of this locus is associated with hypotonia, growth retardation, hyperphagia, mild to moderate learning disability, as well as a range of psychiatric conditions^2, 3^. Upwards of 20% of PWS patients are formally diagnosed with major depressive disorder (MDD), while 50-70% score highly on questionaires for anxious and depressive episodes^4^. Phenotypes of reduced attention and poor impulse control are also highly prevalent and are reported to be intensified in children with PWS compared to their peers with learning disability only^5, 6^. Furthermore, between 16% and 32% of patients experience episodes of psychosis, including delusional ideation and hallucinations^4, 5, 7, 8^.

The prevalence and variation of psychiatric endophenotypes in PWS are influenced by individual genotype. Individuals with PWS can be roughly divided into two groups based on distinct genotype. The first and most prevalent group (∼75%) consists of those that carry a paternally derived deletion spanning the genes of the 15q11-q13 (delPWS). In recent years, several case studies have reported paternally derived microdeletions of the 15q11.2-q13 non-coding RNAs in individuals that exhibit growth retardation, hyperphagia and hypotonia^9-11^. Since these are regarded as the core features of PWS, the region covered by the microdeletions, in which non-coding PEGs *SNORD116* and *IPW* are deleted, is referred to as the PWS critical interval (PWS-cr).

The second most common PWS genotype is maternal uniparental disomy of chromosome 15 (mUPD15), found in about 25% of PWS patients. As well as the core phenotypes, population studies of individuals with PWS report increased incidence and severity of psychosis in those with mUPD15 compared to delPWS, including a higher rate of diagnoses of schizoaffective and bipolar disorders^12, 13^. mUPD15 leads to loss of expression of all PEGs, including those in the PWS-cr, and also to an overexpression of the two maternally expressed genes (MEGs) in the 15q11-q13 imprinted interval^1^. A similar molecular profile (loss of PEGs, overexpression of MEGs) is seen in the rare imprinting centre (IC) mutation genotype (<5%)^14^, which also shows incidence of psychotic illness^15^. These findings have led to the suggestion that the overexpression of MEGs, and specifically *UBE3A*, might be the main contributors to psychiatric illness in PWS^1^, an idea supported by the increased incidence of psychotic illness in individuals carrying maternally-derived copy number variation (CNV) duplications at 15q11-q13^16^. Nevertheless, delPWS patients can also exhibit depression, anxiety and, albeit with reduced incidence and severity, psychotic episodes^7, 17^, and recent genomic studies linked variation in the PWS PEGs *MAGEL2, NECDIN* and *SNORD116*, to scoring highly on the Schizotypal Personality Questionnaire^18, 19^. Taken together, this suggests a potential combinatorial role for both the PWS PEGs and MEGs in psychiatric illness. However, as yet the mechanism leading to psychiatric illness remain unclear.

To address this, we have used mouse models to investigate behavioural and cognitive phenotypes of relevance to psychiatric illness seen in PWS. Previously, we have examined behavioural and cognitive phenotypes in a PWS imprinting centre deletion (PWS-IC) mouse, which models the rare IC mutation genotype and demonstrates loss of expression of PEGs and overexpression of MEGs^14, 20^, matching that observed in individuals with mUPD15. We have shown that the PWS-IC model recapitulates many of the behavioural and cognitive deficits of relevance to PWS, including hypoactivity, abnormal sensory-motor gating, reduced attention and greater impulsivity^20-22^. Here, we examine the behaviour of another PWS mouse model carrying a deletion of PWS-cr, in order to investigate whether the PWS-cr contributes to the endophenotypes of relevance to psychiatric illness and cognition observed in the PWS-IC mouse model. Both models show the growth^20, 23, 24^, endocrine^23, 25^ and hyperphagia^26-28^ phenotypes central to PWS. We tested whether hypoactivity, abnormal sensory-motor gating, reduced attention and greater impulsivity are indeed limited to the PWS-IC mice by conducting the same behavioural characterisation in PWS-cr mice. Furthermore, we attempted to examine the broader molecular changes that may underpin these cognitive and psychiatric endophenotypes specifically by using RNA-seq to examine gene expression changes in the brain of both PWS models. The behavioural differences between models were reflected in a distinct profile of differentially expressed genes (DEGs) and isoform changes between PWS-IC and PWS-cr brain. Critically, the DEGs and isoform changes in PWS-IC, but not PWS-cr, were enriched for genetic variants associated psychotic illness, but interestingly not schizophrenia. Our findings shed light on a molecular basis of behavioural and cognitive problems seen in PWS and may have implications for the treatment of psychotic illness in individuals with genetic lesions at 15q11-q13.

## MATERIALS AND METHODS

### Behavioural experiments

#### Animal husbandry

Due to the genomic imprinting of the 15q11.2-q13 locus, heterozygous mice carrying the deletion on the paternally inherited chromosome are expected to have a complete loss of expression of the critical interval. PWS-cr^m+/p-^ mice (B6(Cg)-Snord116^tm1Uta^/J) were purchased from The Jackson Laboratory. The females were mated with wild type C57BL/6 males in order to generate PWS-cr^m-/p+^ mice, which carry the deletion on the silenced maternal copy of the critical interval mutation and are therefore expected not to exhibit any of the phenotypes associated with it. The behavioural cohort of 24 PWS-cr ^m+/p-^ (16 female, 8 male) and 31 wild type (17 female, 14 male) was produced by breeding PWS-cr^m-/p+^ males with wild type CD1 females in order to exactly replicate the genetic background that we have used in our previous study examining the behaviour of a PWS-IC mouse model^20^. Weaning took place at approximately four weeks of age, when animals were housed in single sex and mixed genotype cages of 2-4 individuals. Ear tissue was collected for genotyping and identification. Experimental testing began at approximately 8-9 weeks of age and took place in the order in which the tests are described below.

The animals were handled daily for two weeks prior to the start of experimentation in order to adapt them to being picked up. All mice were kept at a 12 h light/dark cycle, with the lights being switched on at 7:00 h every morning. Access to water and food was *ad libitum*, until two weeks before the start of the reward preference test, when the animals were gradually introduced to a water deprivation regime. During the first two days of this regime, the animals were given access to water for 4 hours per day. After that, the access to water was limited to 2 hours per day. The water deprivation regime continued all through the reward preference test and the 5-choice serial reaction time task. Water bottles were provided in the home-cages immediately after experimentation.

#### Elevated plus maze and open field

The elevated plus maze (EPM) and open field test (OF) were used to measure anxiety-related behaviours^29, 30^. The EPM consisted of a plus shaped platform, with two enclosed arms (19×8×15 cm) and two open, exposed arms (19×8 cm). The maze was lifted 50 cm above the ground. The animals were placed in one of the enclosed arms of the maze at the beginning of the task, and allowed to freely explore for 5 min. The OF apparatus consisted of a square arena (75×75 cm) enclosed by walls (45 cm). The surface of the arena was divided into central zone (45×45 cm) and outer zone. The animals were placed into the outer zone of the arena and allowed to freely explore for 10 min. The movement of the animals in the EPM and OF was detected by a camera attached to a computer with Ethovision software (XT, Noldus, Netherlands). Time spent in movement during the trials was calculated post-hoc by dividing the distance by the velocity.

#### Locomotor activity

Spontaneous locomotor activity (LMA) of mice was explicitly measured in custom made chambers (21×36×20 cm) fitted with infrared beams, situated 3 cm from either end of the box and 1 cm from the floor of the box. The test was run in complete darkness in order to remove the anxiogenic effect of light. The mice were placed in the boxes and allowed to roam freely for 2 h, while the disturbance of the beams by their movement was recorded by an Acorn computer with an Arachnid software (Cambridge Cognition LTD). Number of beam breaks and the consecutive breaking of the two infrared beams (referred to as a “run”) were considered the main measures of locomotor activity. The animals were assessed at the same time for two consecutive days, in order to assess habituation to a novel environment within and between sessions.

#### Acoustic startle and pre-pulse inhibition

Acoustic startle response (ASR) and pre-pulse inhibition (PPI) were used to test for phenotypes associated with psychotic and affective illness^31^. ASR and PPI were measured in a soundproofed SR LAB startle chamber with a speaker (San Diego Instruments), in which the mice were securely placed inside a plexiglass tube (3.5 cm dia). The trial commenced with 5 min of 70 dB white noise in order to allow for habituation to the environment. After habituation, the mice were exposed to a range of acoustic stimuli, including pulse-alone stimuli at 120 dB and 105 dB above background noise (70dB) for 40 ms, then startle stimuli preceded by 20 ms pre-pulses of 4 dB, 8 dB and 16 dB, 80 ms prior to startle stimuli. All startle and pre-pulse stimuli were presented against a background noise of 70 dB. The startle movement in response to the noise was recorded with a piezoelectric accelerometer and normalized by body weight before analysis. PPI of the ASR was calculated as the percentage reduction in startle amplitude between responses to startle stimuli alone (averaged cross 13 trials) and pre-pulse trials (averaged for the 5 presentations at each pre-pulse amplitude.

#### Reward preference test

Prior to the 5-CSRTT and after a transition to a restricted home-cage water access, animals were gradually exposed to the substance (10% condensed milk in tap water) which was to be used as a reward for the tasks later on. The animals were placed individually in a cage without sawdust, for a 10 min session each day, over a 7-day period. Each cage contained two small plastic containers (∼1 cm high x ∼2 cm diameter) secured to the floor by Velcro, placed equidistant from the end wall, and evenly spaced across the width of the cage. On the first two days, both containers contained tap water. On the following four days, one container contained the condensed milk dilution and the other - water, with the positions alternating each day of testing. On the final day, both containers were filled with the condensed milk dilution. All consumption was measured in grams by weighing the containers before and after the task. The preference to the reward was calculated for days 3 to 6 as a percentage of total volume consumed.

#### 5-choice serial reaction time task

The 5-choice serial reaction time task (5-CSRTT) is commonly used to measure visuo-spatial attention and impulsivity in rodent models of disease^32, 33^. The 5-CSRTT was commenced on the day following the final day of the reward preference test and broadly followed the protocol described by^34^. For this task, mice were trained to respond to a brief visual stimulus appearing pseudo-randomly at one of five locations, in order to gain a condensed milk reward. The variables analysed as indicative of performance were accuracy (ratio of correct to total trials), omissions (ratio of omitted trials to total trials), and premature responses (number of responses that took place before the stimulus was presented). Other variables such as completed trials (calculated as ratio out of 60), correct reaction time, reward collection latency, number of food magazine entries and total number of trial perseveration were also kept track of for further analysis. Subjects were trained to a set baseline performance observed at a stimulus duration of 0.8 s (>30 completed trials, >75% accuracy, and <25% omissions). Full details can be found in Supplementary information.

The conditions of the experiment were manipulated in order to study various aspects of attention and impulsivity. One of the manipulations entailed shortening the stimulus duration (SSD). During this session, stimuli of 0.6 s, 0.4 s and 0.2 s were interspersed in pseudorandom order along with the baseline stimulus duration of 0.8 s, in order to increase the difficulty of the task and to affect the accuracy of performance. In another manipulation lengthened it is (LITI) of 7 s, 9 s and 11 s were interspersed in pseudorandom order along with the baseline inter-trial interval of 5 s, in order to induce premature responses and test for impulsive behaviour.

#### Data analysis

All data were analysed using R Studio 1.1.383 (R Studio, Inc). Linear models (lm) and generalized linear models (glm) were built with genotype and sex as fixed effects. Since interactions between sex and genotype were not observed, the effect of sex is not reported in this paper. However, the data, which are freely available on the ‘Open science framework’ (https://osf.io/wnx8r/) are identifiable and therefore separable by sex. Logistic glm from the binomial family was used to analyse all proportional data. Mixed lms and glms with mouse ID as a random effect were built using the lme4 package^35^, for data in which multiple measures were taken from each individual at different conditions. In the LMA mixed model, day of testing and time bins were added as fixed effects. In the ASR mixed model, startle trial number was added as fixed effect. In the PPI mixed model, the different pre-pulse decibels (4 dB, 8dB, 16 dB) were added as fixed effects. For the 5-CSRTT SSD and LITI manipulations, stimulus duration and inter-trial interval were also added as fixed effects. The p-values and chi-square values (χ^2^) presented in the results were calculated using likelihood ratio tests run with the inbuilt ANOVA function of R and comparing the full statistical models to reduced models with genotype removed as a fixed factor.

### RNA sequencing

#### Tissue collection, preparation, and RNA sequencing

For the RNA sequencing experiments, whole brain neonatal tissue was taken from 6 PWS-cr mice and 6 of their wild type littermates (from 4 separate litters), and from 4 PWS-IC^m+/p-^ mice and 6 of their wild type littermates (from 2 separate litters). Tissue was collected first thing in the morning (9-10:00am) following birth and was snap frozen on dry ice and stored at -80°C until. RNA was extracted using the Direct-zol RNA Miniprep kit. Libraries were prepared with the KAPA mRNA HyperPrep kit (Roche Sequencing solutions) from total RNA of RQN > 8.5 as assessed by the Fragment Analyser system (Agilent Technologies). 800 ng of polyA selected RNA was used for each sample for a paired and unstranded sequencing approach. The target read depth was 80 million reads per sample. The mRNA was fragmented at 94°C for 6 minutes to achieve a mean insert size of 200-300 bp. 8 cycles were used in the final amplification. A final bead clean-up step was added at the end of the protocol before the libraries were taken forward for NGS on an Illumina HiSeq4000 system.

#### RNA sequencing analysis

For analysis of differential gene expression, sequencing reads were mapped to the mouse genome (version mm10) using the STAR software package^36^. Unmapped reads were output into the BAM file with the Within KeepPairs option, which records unmapped mates for each alignment and keeps any unsorted output adjacent to its mapped mate. Multimapping alignment was done with a random order. The quantification of annotations was calculated with the GeneCounts option which outputs count reads per gene. The featureCounts software^37^ was used to count the raw reads, which were subsequently analysed in R Studio 1.1.383 using the DESeq2 package^38^, with genotype and litter as fixed effects into the statistical model. Hierarchical clustering was performed with the heatmap.2 package in R, which uses Euclidean distances to calculate difference between data points. The clustering showed an outlier in both PWS-cr and PWS-IC samples, which were removed from further analysis. The UCSC genome browser was used for visualization of the data. Differential isoform usage was pseudo-aligned to the mouse transcriptome (mm10) with the Kallisto software ^39^ which was also used to quantify the raw reads. The Kallisto files were then analysed in R studio via DEXSeq with the IsoformSwitchAnalyzeR package, which measures isoform usage by quantifying the fraction of isoform expression from the parent gene expression. The difference in usage between isoforms is then used to measure the effect size^40-42^. Prediction of protein domains was run on EBI’s Pfam webserver, which performs biosequence analysis using hidden Markov models^43^. The Pfam data were then integrated back into IsoformSwitchAnalyzer for final statistical analysis. Gender and litter were added as covariates into the statistical model. All p-values were adjusted using the Benjamini-Hochberg method. The data from the PWS-cr and PWS-IC models was analysed separately since difference between the wild type littermates of the two groups did not allow for pooling of the data. Genes that were differentially expressed or exhibited differential usage of isoforms between genotypes (padj<0.05) were pooled together for functional enrichment analysis using the g:Profiler web server ^44^.

#### Common variant enrichment analysis

Genes that were differentially expressed or exhibited differential usage of isotopes between genotypes (padj<0.05) were pooled together for common variant enrichment analysis. The mouse gene IDs were converted to their homologous human gene IDs. The genes were run for gene set analysis through MAGMA along with GWAS summary statistics for schizophrenia, psychosis, and chronic kidney disease ^45-47^. Gene locations build 37 and reference data from European population was downloaded from CNCR CTGLab (https://ctg.cncr.nl/software/magma).

## RESULTS

### Elevated plus maze and open field test

As expected, on the EPM all mice spent significantly more time of the 5 min session in the closed arms of the maze compared to the more anxiogenic open arms of the maze (χ^2^ _(1)_=138.98, p<0.001), but no effect of genotype on percentage of time spent in the open arms of the EPM (F_(3,50)_=0.132; p=0.361) (Figure 1A), or on distance travelled through the area of the maze (F_(3, 50)_=2.352, p=0.174). Number of head dips and stretch attends were also recorded as a measure of explorative behaviour through the EPM and showed no significant differences between PWS-cr mice and their wild type littermates (F_(3, 50)_=0.7832, p=0.989; F_(3, 50)_=1.933, p=0.151).

**Figure 1.**
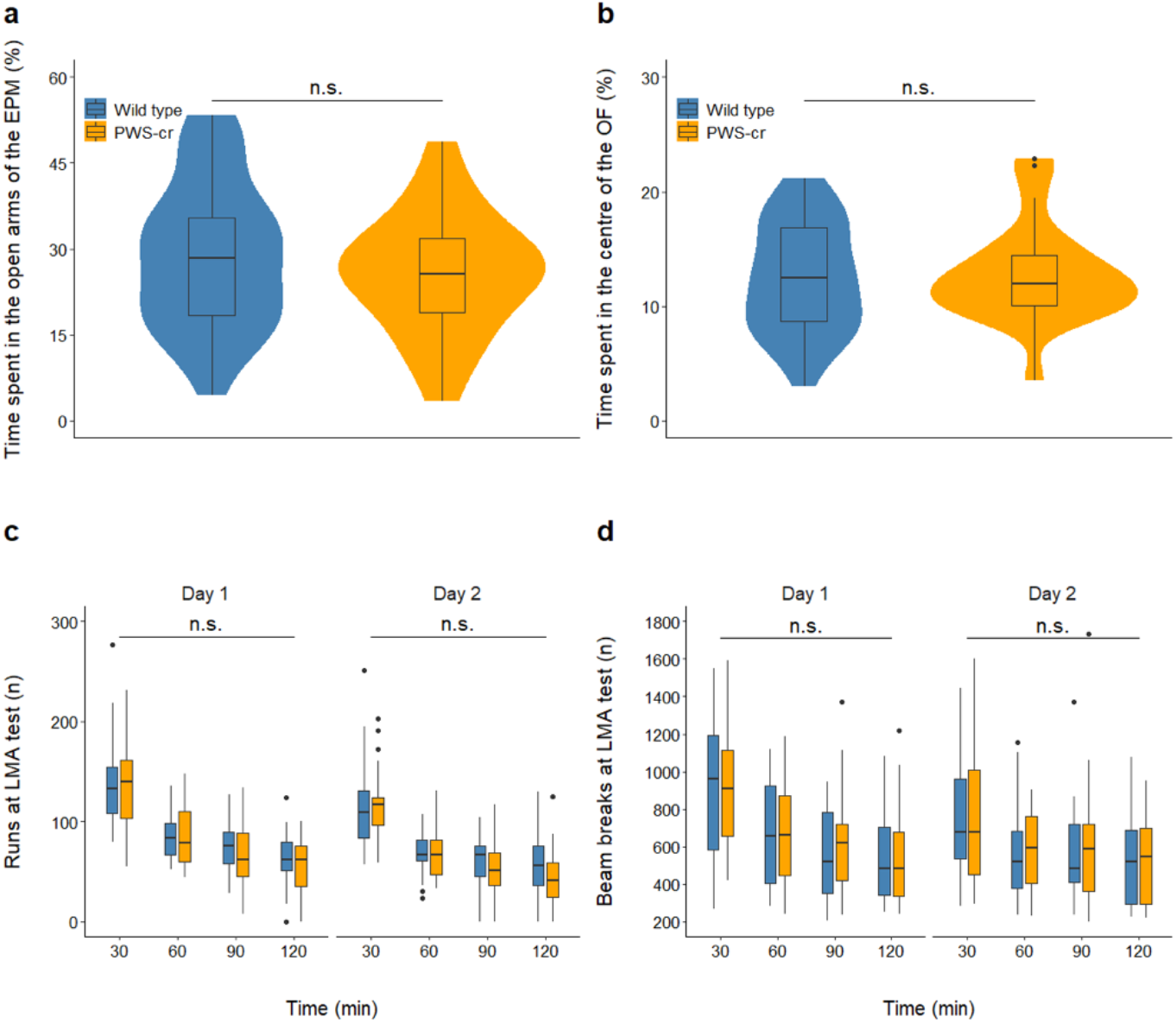
Behaviours recorded at the elevated plus maze, open field arena and locomotor activity test. The elevated plus maze task (a) showed no effect of genotype on percentage of time spent in the open arms of the maze (PWS-cr=23, WT=31). Similarly, data from the open field test (b) showed no difference in percentage of time spent in the centre square of the arena between PWS-cr mice (n=23) and their wild type littermates (n=31). Locomotor activity was tested in custom made chambers with fitted infrared beams, which tracked the movement of the animals in two-hour sessions on two consecutive days. Number of beam breaks (d) and number of runs from one end of the chamber to the other (c) were considered the main measures of activity. The test showed decreased locomotor activity between and within sessions (p<0.001), which is indicative of habituation to the environment. There was no effect of genotype on the behaviour examined by this test (PWS-cr=23, WT=31).

During the OF test, mice spent significantly more time of the 10 min session in the outer areas of the arena than in the more anxiogenic centre square χ^2^ _(1)_=452.2, p<0.001), but there was no effect of genotype on percentage of time spent in the centre square (F_(3, 50)_=0.084; p=0.361) (Figure 1B) or on distance travelled through the arena (F_(3, 50)_=4.55; p=0.472). Overall, the data collected from the EPM and OF show no differences in behaviour between PWS-cr mice and their wild type littermates. Our results suggest no obvious indications of anxiety phenotype in the PWS-cr mouse model.

### Locomotor activity

There was no effect of genotype on total number of beam breaks made in the LMA test over the span of the two sessions χ^2^ _(2)_=4.625, p=0.099), or on total number of runs made from one end of the chamber to the other χ^2^ _(2)_=1.237, p=0.539). Further analysis of the data in 30 min time bins showed that all animals exhibited habituation to the environment within and between sessions, as indicated by the steady decrease of number of runs (Figure 1C; χ^2^ _(3)_=347.75, p<0.001; χ^2^ _(1)_=49.953, p<0.001) and beam breaks (Figure 1D; χ^2^ _(3)_=197.83, p<0.001; χ^2^ _(1)_=33.562, p<0.001) over time. However, there was no statistically significant difference in habituation behaviour between the two genotypes χ^2^_(2)_=3.342, p=0.342; χ^2^ _(2)_=1.761, p=0.222). Overall, the results from this test showed no locomotor activity phenotype in the PWS-cr mice.

### Acoustic startle response and pre-pulse inhibition

Acoustic startle response (ASR) and pre-pulse inhibition (PPI) to acoustic startle response were measured at pulse intensities of 120 dB and 105 dB. A total of thirteen ASR-alone pulses were presented to the mice at the two intensities, and both induced a significantly reduced startle response in the PWS-cr mice (Figure 2A; χ2 _(2)_=6.5292, p=0.03821; Figure 2B; χ2 _(2)_=14.562, p<0.001). Pre-pulses of 8 dB and 16 dB played 70 ms prior to the 120 dB and 105 dB pulses inhibited the startle response up to mean average of 74.09% (±1.84% SEM) and 56.1% (±4.86% SEM) respectively, indicating operational sensory-motor gating in both groups of mice (Figure 2C, 2D). Genotype had no effect on pre-pulse inhibition at either pulse intensity (χ2 _(2)_=5.376, p=0.251; χ2 _(2)_=2.556, p=0.635). Overall, the results from the ASR and PPI tests show no sensory-motor gating phenotypes in the PWS-cr mouse model, but a reduced startle response to acoustic stimuli.

**Figure 2.**
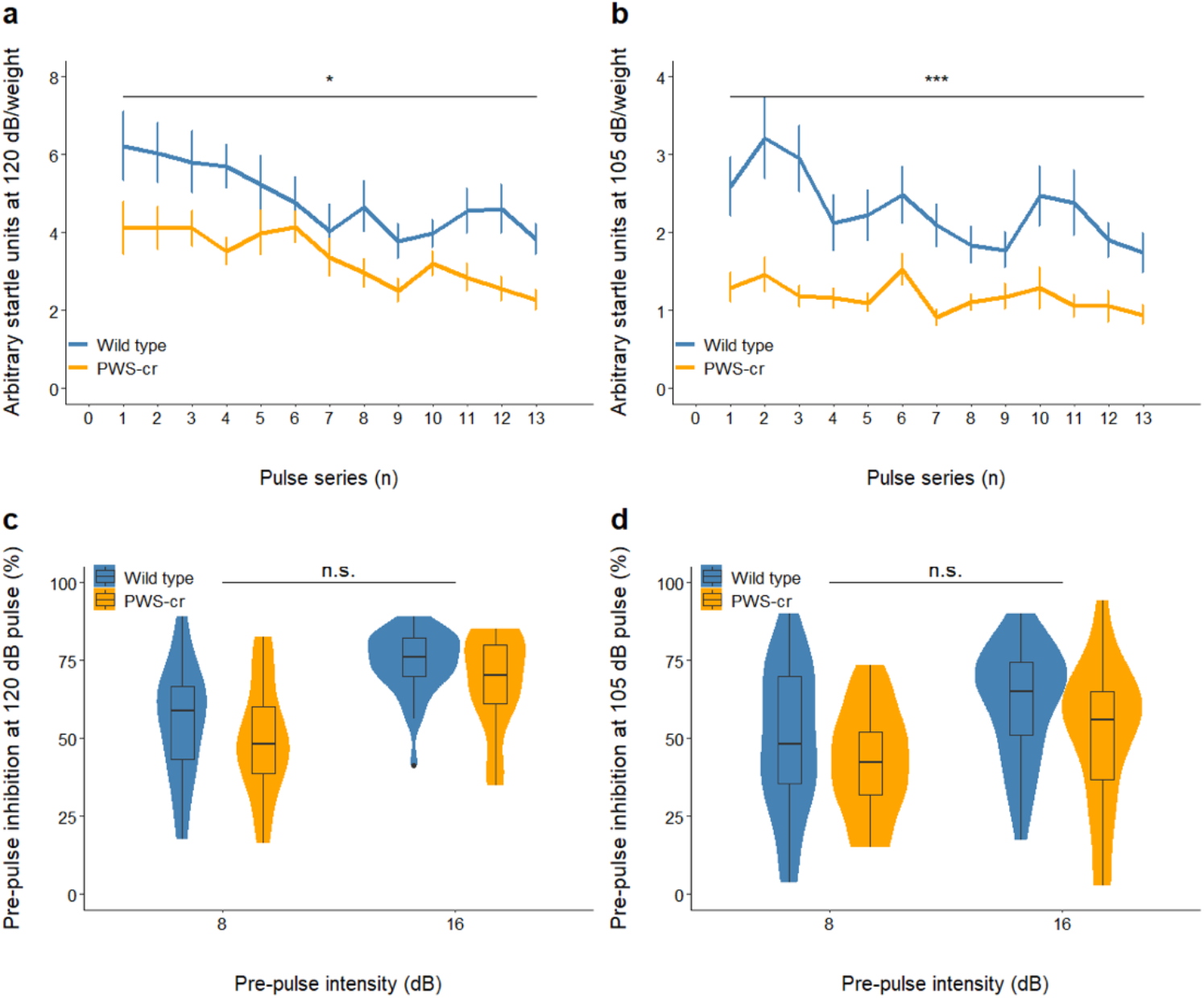
Acoustic startle and pre-pulse inhibition. The startle response to a series of 13 acoustic stimuli at 120 dB (a) and 105 dB (b) was significantly reduced in PWS-cr mice (n=23) compared to their wild type littermates (n=31) (* indicates p<0.05, *** indicates p<0.001). Pre-pulse inhibition of the acoustic startle response was measured by presenting noises of 8 dB and 16 dB 70 ms before the pulses of 120 dB (c) and 105 (d). The pre-pulse inhibition data showed operational sensory motor gating, which was not affected by genotype (PWS-cr=23, WT=31).

Following the 120dB and 105dB ASR and PPI schedules, mice were exposed to a gradual increasing ASR-alone pulse intensity, ranging from 70dB to 120dB. As expected, the increasing intensity resulted in increasing response (χ^2^_(12)_=253.28, p<0.001). These data suggest that the difference in ASR between PWS-cr and wild type mice emerges at >100dB intensity (Supplementary Figure 1). However, there was no effect of genotype (χ^2^_(12)_=20.49, p=0.058) and no interaction between genotype and pulse intensity (χ^2^_(10)_=9.27, p=0.507).

### 5-choice serial reaction time task

Behaviours relating to visuo-spatial attention and control response were investigated in the 5-CSRTT, where mice were trained to respond correctly to stimuli in order to receive condensed milk dilution as a reward. Prior examination showed an increased preference towards condensed milk dilution over water in all animals (χ^2^ _(3)_=68.461, p<0.001), thus validating its use as a motivational reward.

There was no significant distinction between the preference to the reward between PWS-cr and wild type (Figure 3A; χ^2^ _(2)_=0.5827, p=0.7473) and thus no indications of increased interest in the PWS-cr mice towards the reward, which was an important consideration in the interpretation of the results from the operant tasks.

**Figure 3.**
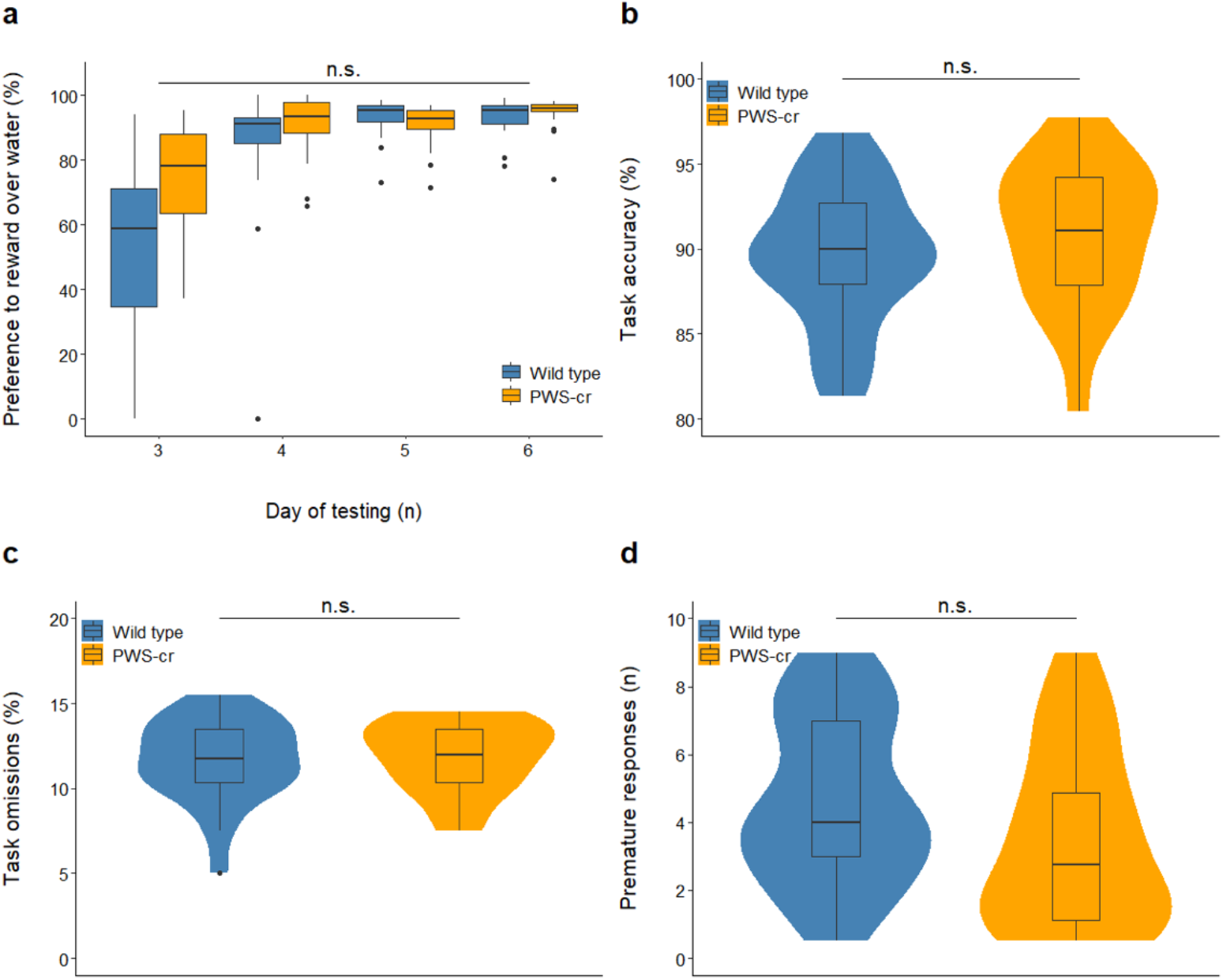
Condensed milk test and 5-CSRTT performance at baseline conditions. Preference to the condense milk reward over water was tested over the span of days 3 to 6 from the condense milk test (a). Over the course of these four days the preference to the reward increased significantly (p<0.001) as expected for the purposes of the 5-CSRTT, and this tendency was not affected by genotype (PWS-cr=21, WT=22). Performance at baseline conditions of the 5-CSRTT also showed no difference between PWS-cr mice (n=20) and their wild type littermates (n=23) in accuracy of performance (b), percentage omissions (b) and number of premature responses (b).

The average number of sessions required to reach baseline performance in the 5-CSRTT did not differ between PWS-cr (mean 98.6, ±7.3 SEM) and wild type (mean 90.8, ±5.7 SEM) mice (F_3,40_=0.703, p=0.802). Performance at baseline conditions of the 5-CSRTT showed no effect of genotype on task accuracy (Figure 3B; F_(3, 42)_=0.127, p=0.724), omissions (Figure 3C, F_(3, 42)_=0.104; p=0.749), premature responses (Figure 3D; F_(3, 40)_=0.515, p=0.361) and all other parameters recorded at these conditions of the task.

The conditions of the 5-CSRTT were manipulated in order to tease out subtler aspects of attention and impulsivity. Behaviour was inspected at stimulus duration of shortened presentation, all introduced in pseudorandom order within one 5-CSRTT session. The shortened stimulus duration task lowered the overall task accuracy (χ^2^ _(3)_=41.114, p<0.001), and this performance decline was not influenced by genotype (Figure 4A; χ^2^ _(1)_=1.642, p=0.2). There was no effect of genotype on percentage omissions (Figure 4B; χ^2^ _(1)_= 0.357, p=0.551) or any of the other parameters that were investigated in the task.

**Figure 4.**
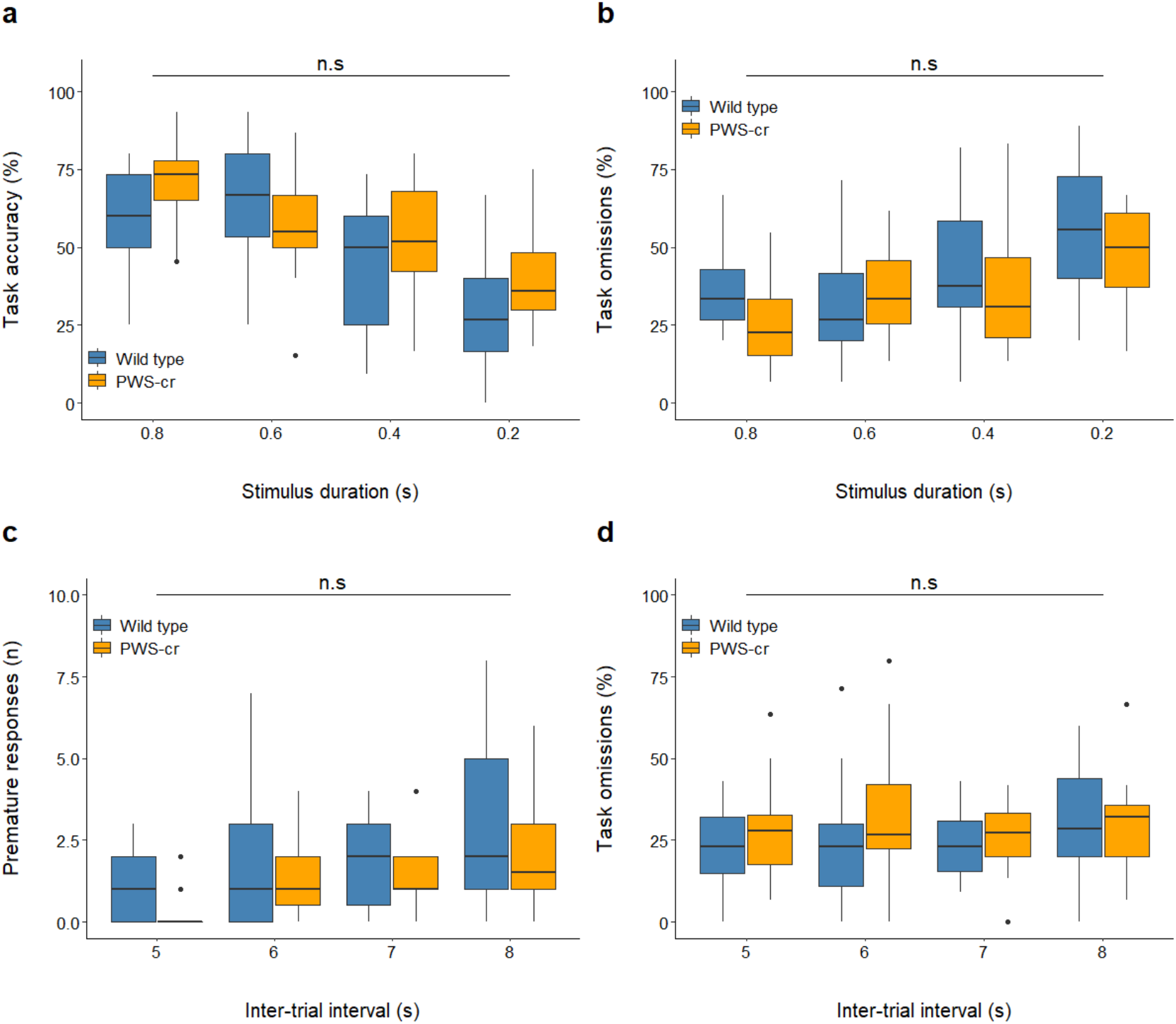
Shortened stimulus duration and increased inter-trial intervals at the 5-CSRTT. Shortening the stimulus duration at the 5-CSRTT reduced performance accuracy (p<0.001) (a) and increased percentage of trial omissions (p<0.001) (b) but revealed no difference between the behaviour of PWS-cr mice (n=15) and their wild type littermates (n=21). Increasing the inter-trial intervals lead to an increase in premature responses (p<0.001), which was not affected by genotype (PWS-cr=18, WT=24) (c). Omissions were also not affected by genotype in this task manipulation (d).

An increase of the inter-trial intervals was introduced in a different session in order to induce premature responses in the task and tease out a potential impulsivity phenotype. The inter-trial interval increase lead to a rise in number of premature responses as predicted (χ^2^ _(3)_=32.131, p<0.001), but this tendency was not affected by genotype (Figure 4C; χ^2^ _(2)_= 4.966, p=0.084). The percentage of omissions at this task was also not significantly different between PWS-cr mice and their wild type littermates (Figure 4D; χ^2^ _(2)_= 0.015, p=0.992), and neither were any of the other examined parameters. Overall, the data collected from the 5-CSRTT showed no indications of an attention or impulsivity phenotype in the PWS-cr mouse model.

### RNA-sequencing of whole brain neonatal tissue

The results from our behavioural study demonstrated a vastly different behavioural and cognitive profile in the PWS-cr mice compared to the previously examined PWS-IC mouse (Table 1). In order to investigate the gene expression changes underlying these behavioural differences, we conducted an RNA-sequencing assay on whole brain tissue from both mouse models in parallel. While all behavioural tasks were conducted on adult individuals, for the RNA-sequencing study we took tissue from neonatal mice, as genetic risk for psychiatric illness has been shown to manifest from early developmental stages^48, 49^. Furthermore, qPCR assay of PWS genes in wild-type and PWS-IC mouse whole brain tissue at three developmental stages (E13.5, E18.5, P0), showed that *Snord115, Snord116, Necdin* had significantly reduced expression in the PWS-IC samples, while *Ube3a* was significantly overexpressed (Supplementary figure 2; t(6) = 11.788, p < 0.001; t(6) = -6.415, p < 0.001; t(6) = -24.033, p < 0.001; t(6) = -12.956, p < 0.001), as well as at stage P0 (t(8) = -3.573, p = 0.007; t(8) = -6.297, p < 0.001; t(8) = -7.4468, p < 0.001; t(8) = -6.415, p = 0.033).

**Table 1.** Summary table comparing behavioural and cognitive findings in the PWS-IC and PWS-cr mouse model for Prader-Willi syndrome. The PWS-IC mice exhibit a range of behavioural phenotypes of relevance to psychiatric illness, in particular increased startle response and decreased PPI and reduced attention^17, 18^. Testing the PWS-cr mice in equivalent conditions (same genetic background, same behavioural equipment, testing rooms, and protocols) revealed no differences, apart from a reduction in acoustic startle response.

| Behavioural domain and measures |  | Behaviour in PWS model compared to WT littermates |  |
| --- | --- | --- | --- |
|  |  | PWS-IC<sup>17,18</sup> | PWS-cr |
| Anxiety | EPM % open arm | ≡ | ≡ |
|  | OF % centre | ≡ | ≡ |
| Activity |  | ↓ | ≡ |
| Sensory motor gating | Acoustic startle response | ↑ | ↓ |
|  | PPI of startle response | ↓ | ≡ |
| Attention (5-CSRTT) | Accuracy | ↓ | ≡ |
|  | Omissions | ↑ | ≡ |
| Impulse control (5-CSRTT, baseline) |  | ≡ | ≡ |

#### PWS-cr differentially expressed genes and isoforms

Visualization of the aligned RNA-sequencing reads in the UCSC browser confirmed that the deletion in the PWS-cr mouse model mode spans all copies of *Snord116* snoRNA and five of the six *Ipw* exons (Supplementary Figure 3). However, differential gene expression analysis showed only very subtle differences between the PWS-cr mice and their wild type littermates after Benjamini-Hochberg adjustment for multiple testing. In total, there were seven differentially expressed genes (DEGs) in brain tissue from the PWS-cr mouse model (Figure 5a and Appendix 3). Among them was the *Snhg14* non-coding RNA, which acts as a host for the critical interval (z=21.846, padj<0.001). The *Mafa* gene encoding a transcription factor that regulates pancreatic beta cell-specific expression of the insulin gene was upregulated in the absence of the critical interval (z=-7.014, padj<0.001). Notably, the *Necdin* growth suppressor which is one of the imprinted genes of the PWS locus, was upregulated in the PWS-cr mouse brain tissue (z=-8.621, padj<0.001). The remaining four DEGs were predicted genes of unknown function, two of which (*Gm44559* and *Gm44562*) fall within intron 4/12 of the *Ube3A*, and *Gm44831* which falls into *Ipw* in the critical interval.

**Figure 5.**
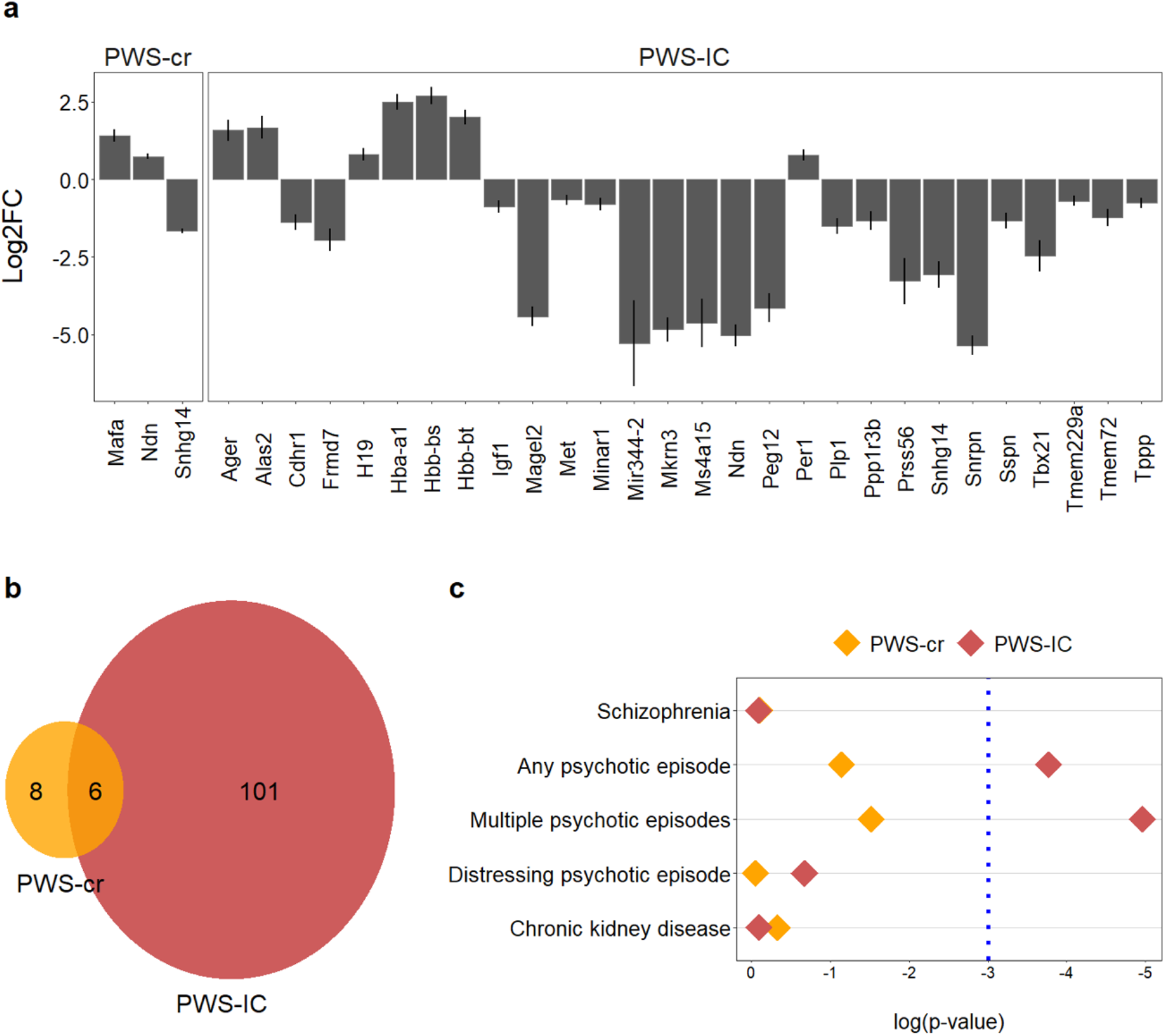
RNA-sequencing assay. Differential gene expression analysis (a) showed an overexpression of *Necdin* and *Mafa* in the PWS-cr tissue, as well as an expected loss of expression of the long non-coding RNA *Snhg14* that acts as a host for the critical interval. Among the differentially expressed genes in the PWS-IC tissue were the genes from the PWS locus, and notably the circadian clock regulator *Per1* and the insulin growth factor *Igf1*. Predicted genes of unknown function were excluded from the graph. The y axis on this graph (a) shows log2 fold change of expression, error bars indicate standard error of the mean. Differentially expressed genes and isoforms were pooled for enrichment analyses; PWS-cr and PWS-IC pooled genes had 6 genes in common (b). Gene set analysis of pooled differentially expressed genes and isoforms (c) showed a significant enrichment of common genetic variants associated with the experience of psychotic episodes, but not schizophrenia. GWAS summary data from a chronic kidney disease study was used as a negative control.

In addition to differential gene expression, the read depth used in our RNA-seq analysis allowed analysis of isoform switches with functional consequences in order to investigate whether the genes in the critical interval play a regulative role in alternate splicing. Seven genes showed significantly different patterns of splicing in the PWS-cr mouse brain (Appendix 4), and there was no overlap between them and the DEGs. Among these was dual-specificity tyrosine-phosphorylation-regulated kinase (*Dyrk3*), which plays a role in the dissolution of stress granules and in the early secretory pathway^50^. The DEGs and isoforms were pooled together for a gene ontology analysis, which revealed no enrichments.

#### PWS-IC differentially expressed genes and isoforms

In contrast to PWS-cr, analysis of PWS-IC mouse brain tissue RNA-sequencing data primarily clustered by genotype and revealed 59 significantly DEGs compared to their wild-type littermates (Figure 5a; Appendix 5). This included *Snhg14*, the non-coding RNA that host the *Snord116* and *Ipw*, but also, as expected, the other PWS genes such as *Snrpn, Necdin, Mkrn3*, and *Magel2*. Outside the PWS cluster there were other notable DEGs, including the circadian clock regulator *Per1* (z=-4.25, padj=0.012), Plp1 (z=6.11, padj<0.001) and insulin growth factor *Igf1* (z=4.164, padj=0.015).

Analysis of isoform switches identified 48 differentially spliced genes, and there was no overlap with the differentially expressed genes (Appendix 4). Differentially expressed genes and isoforms were pooled together for a gene ontology term analysis, which were enriched for molecular functions, cellular compartment, and biological compartments relevant to oxygen transport (Appendix 6).

Comparing the combined DEG and isoform data for the two models directly shows that there were a greater number of differences in the PWS-IC mice overall (Figure 5b). As expected, a good proportion (∼50%) of the changes seen in PWS-cr were shared with those found in PWS-IC (Figure 5b). However, this overlap was limited to DEGs, as there were no common differentially spliced genes in the PWS-cr and PWS-IC samples.

#### Enrichment of common genetic variants

We pooled the differentially expressed and differentially spliced gene sets (padj<0.05) together for each mouse model in order to look for enrichment of common genetic variants associated with schizophrenia or psychotic episodes. The analysis was performed using MAGMA’s regression gene-set computation with the datasets from genome wide association studies (GWAS) that utilize the UK Biobank, ClozUK and CKDgen consortium databases respectively^45-47^. The results showed no enrichment of genetic variants of interest in the PWS-cr samples (Figure 5c). In contrast, results from the PWS-IC model indicated an enrichment of SNPs common for the experience of ‘any’ and ‘multiple’ psychotic episodes (Appendix 7; β=0.215, p=0.023, β=0.271, p=0.007), but interestingly not for schizophrenia (Figure 5c; β=-0.206, p=0.913). As expected, neither gene set was enriched for the negative control dataset, common genetic variants associated with chronic kidney disease (β=-0.161, p=0.909).

## DISCUSSION

Our aim in this study was to investigate the differential contribution of genes in the Prader-Willi syndrome imprinted interval to psychiatric illness using mouse models. We have previously demonstrated abnormalities in a number of behavioural and cognitive endophenotypes of relevance to psychiatric illness in a PWS imprinting centre deletion (PWS-IC) mouse model. Here we show that these same behavioural and cognitive measures are not altered in a mouse model for PWS that has a deletion limited to the PWS critical interval (PWS-cr mice). This suggests that the loss of expression of the PWS-cr genes does not contribute to the behaviours previously observed in PWS-IC mice. We then used RNA-seq to examine the molecular bases of these differences in behaviour between the PWS-IC and PWS-cr models. Although a number of interesting gene expression and alternate splicing events were detected in the brains of PWS-cr mice, a greater number were seen in the PWS-IC model. Moreover, these changes in gene expression and splicing in the PWS-IC brain were enriched for common genetic variants associated with the experience of psychotic episodes, but not those associated with schizophrenia. These findings shed light on the nature of psychotic illness in PWS and have implications for possible therapeutic strategies.

The behaviour of mice carrying a deletion of the PWS-cr has been extensively characterized with respect to various aspects of the core PWS phenotype including hunger and sleep^23, 51-53^, but behaviours relating to psychiatric illness have not been directly studied. Our previous behavioural studies of the PWS-IC model indicated reduced activity, abnormal sensory-motor gating, and deficits in attention^20^. In contrast, here our studies of the PWS-cr mouse model suggests that deletion of the critical region has only very subtle effects on these same behavioural measures. Specifically, the PWS-cr mice showed no abnormalities in anxiety as measured by the EPM and OF tests, locomotor activity, sensory-motor gating as measured by PPI, or attention and impulsivity as measured by the 5-CSRTT. While we cannot rule out Type II error, we note that the PWS-cr behavioural cohort was considerably larger than that used for the PWS-IC mice. Critically the behavioural analysis of PWS-cr mice was conducted in an identical manner to that previously used for the PWS-IC model, including the equipment, procedures, software, location (both Institute and rooms) and genetic background strain (F1 cross between C57Bl/6 and CD1). Therefore, we are confident the comparison between these behavioural studies is robust and valid.

Of the behaviours assessed only ASR was altered in PWS-cr mice. However, in contrast to the enhanced startle response seen in PWS-IC mice, PWS-cr exhibited a decreased startle response. A reduced acoustic startle response could potentially be confounded by hearing loss, and this cannot be completely ruled out here. However, this explanation would seem unlikely given the sensitivity of the response in PWS-cr mice to different startle pulse intensities (Supplementary figure 1), and that there are no records of hearing loss in any mouse models for PWS, or indeed in PWS patients. The combined phenotype of an increased startle response and deficient pre-pulse inhibition seen in the PWS-IC mouse model is a long established endophenotype seen in psychotic illness^54-56^ that has been repeatedly shown to have translational utility^31^. Consequently, the lack of any PPI deficit in PWS-cr deletion model suggests that their reduced startle response is a distinct behavioural outcome. Reduced startle response has been reported in individuals with major depressive disorder and anhedonia, as well as an effect of anxiolytic drugs^57-59^. With this in mind, the reduced ASR seen in PWS-cr mice could fit the psychiatric profile of individuals with PWS that carry deletions spanning the critical interval, who are prone to depression and anxiety, but not psychosis^9-11^. However, a broader variety of behavioural analysis is required to properly establish a depression phenotype in the PWS-cr mice.

The behavioural comparison of these two models indicates that the PWS-cr mice do not exhibit the same range of cognitive and psychotic illness endophenotypes as seen in the PWS-IC mice (Table 1). We next explored the molecular bases for these behavioural differences using RNA-seq. Analysis of RNA-seq data comparing PWS-cr whole brain with wild-type littermates revealed only very subtle changes of gene expression and alternative splicing, with seven differentially expressed genes (four predicted transcripts) and five differentially spliced transcripts. This may seem at odds with previous findings demonstrating differential expression of upwards of 200 genes in PWS-cr mice^27, 60, 61^. However, these studies looked at gene expression in more focused tissues (e.g. hypothalamus) or cell types, whereas our study examined bulk whole brain, therefore obscuring some of the subtler region- or cell-specific changes and revealing only the strongest effects across the whole brain. In addition to *Snord116* and *Ipw*, that are directly affected by the mutation (Supplementary figure 2), our data suggest that the PWS-cr deletion also leads to mis-expression of the PWS interval gene *Necdin*, which is an important regulator of neuronal outgrowth and differentiation^62, 63^, loss of which is associated with motor deficits and enhanced learning and memory^64, 65^. Whether the increased expression of *Necdin* is due to loss of suppression by *Snord116* and/or *Ipw*, in manner similar to the regulatory loop between *Snord116* and *Snord115*^60^ or the suppressive role of *IPW* on the maternally expressed genes from the *DLK1-DIO3* imprinted cluster^61^, remains to be established. Nevertheless, it could be that this altered expression contributes to the learning and memory deficits seen in PWS-cr mice^66^.

In contrast to the limited changes seen in the PWS-cr model, RNA-seq analysis of PWS-IC whole brain revealed a markedly larger number of DEGs and isoform changes. We suggest these gene expression changes may contribute to behavioural and cognitive deficits seen in the PWS-IC but not the PWS-cr mice and, by extension, may underpin the increased incidence of psychotic illness seen in individuals with imprinting centre deletions or mUPD^13^. In line with this, as well as altered expression of a greater number of genes from within the PWS interval, there were changes in a number of other neurally important genes. These included reduced expression of the myelin protein proteolipid protein *Plp1* which has been linked to spatial attention^67^. There was also a significant increase in expression of the circadian rhythm gene, *Per1*^68^ in PWS-IC brain. Critically, all our samples were collected within a limited time window in the morning, suggesting that any change in expression may be due to loss of expression of several PWS genes with known circadian rhythm effects^52, 69^. Of most direct relevance to psychiatric illness were the changes to the gamma-aminobutyric acid (GABA) A receptor subunit gamma 3 gene *Gabrg3*, which exhibited increased usage of a truncated isoform lacking the neurotransmitter-gated ion-channel ligand binding domain (NGIC-LBD) and neurotransmitter-gated ion-channel transmembrane region (NGIC-TMR). The GABA receptors play a key role in the function of the central nervous system and this alternate isoform usage could have an effect on various phenotypes including psychotic illness^70, 71^. This further ties in with a study by Webb et al.^72^, which showed that in a significant proportion of the PWS-del individuals that exhibit psychosis, carry a larger deletion that spans part of *GABRG3*. This suggests that the GABA receptor could also be contributing to psychotic illness in individuals with the mUPD15 genotype due to reduced functionality caused by differential isoform usage.

In order to more formally assess the overall relevance of these transcriptomic changes to psychiatric illness we examined the DEGs and isoforms for enrichment of genetic variants relevant to psychosis. The GWAS data we used to look for genetic correlation between our samples with the occurrence of psychotic episodes was from a recent UK Biobank study that sampled individuals with a history of psychotic experiences including hallucinations and delusional ideation, but specifically without a diagnosis of schizophrenia^47^. The authors found a shared genetic liability with schizophrenia from an external GWAS dataset^46^, which we also used for our analysis here. Unsurprisingly given the limited number of changes, there was no enrichment in the gene expression changes seen in the PWS-cr neonate brain. However, there was an enrichment of genetic variants common to the experience of psychotic episodes in the gene expression changes seen in the PWS-IC neonate brain. Furthermore, among the genes that carried common SNPs of psychotic experiences were the paternally expressed PWS genes *Magel2, Necdin, and Mkrn3*. The findings substantiate the relevance of the gene expression changes and behavioural endophenotype observed in the PWS-IC mouse model to the psychiatric profile of individuals with individuals with an IC deletion or a mUPD of chromosome 15, but also suggest a contribution of some of the PWS PEGs to these phenotypes. Strikingly, there was not an enrichment of genetic variants associated with schizophrenia in the gene set analysis, which suggests distinct genetic liability of psychosis in PWS from schizophrenia (Figure 6). This finding provides a biological signal that confirms the observation that the psychotic illness seen in PWS is distinct from schizophrenia^12^ and should not be treated in the same manner^17^.

**Figure 6.**
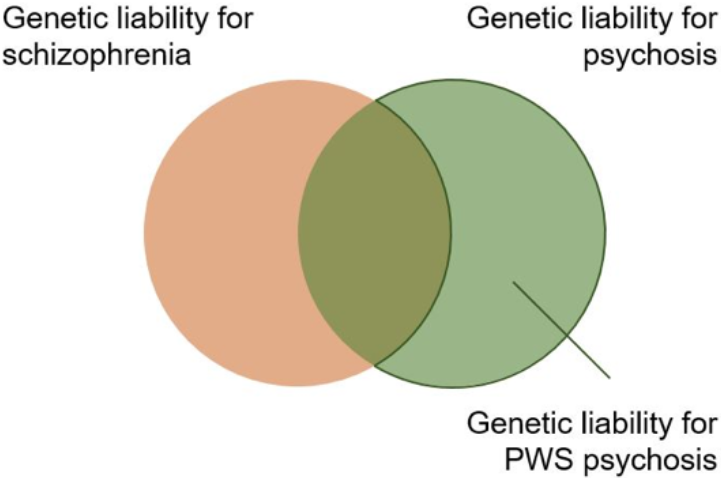
Model of genetic liability for psychosis in individuals with PWS. Gene set analysis of differentially expressed genes and differentially used isoforms in whole neonatal mouse brain showed enrichment of genes relevant to psychotic experiences, but not schizophrenia. Although schizophrenia and psychotic experiences have a shared genetic liability, they are ultimately distinct psychiatric conditions phenotypically and genetically. We propose that the genetic liability for psychosis in PWS is distinct from that of schizophrenia.

Overall, our findings suggest a cumulative effect of several of the genes from the PWS locus on the behavioural endophenotypes typical of Prader-Willi syndrome. The critical interval deletion, albeit important in the core phenotypes seen in PWS models, only induced very subtle behavioural and cognitive differences in the studies we conducted. By comparing the gene expression and isoform profile of the PWS-cr and PWS-IC lines were able to identify brain molecular changes of relevance to the psychiatric profile of PWS, underlined by the enrichment of genetic variants for psychotic episodes in the PWS-IC transcriptomic changes. The fact that genetic variants for schizophrenia showed no enrichment reflects the distinct nature of psychotic illness in PWS and could have implications for therapeutic strategies used to treat psychosis in individuals with PWS.

## DATA AVAILABILITY

All behavioural data (Appendices 1 and 2) and all RNA-seq data, including gene ontology and genetic enrichment data (Appendices 3 to 7) are publicly available at the following ‘Open science framework’ link: https://osf.io/wnx8r/

