## Supplementary for "Comparison of mouse models reveals a molecular distinction between psychotic illness in PWS and schizophrenia"

**SUPPLEMENTARY MATERIALS AND METHODS**

***Detailed 5-choice serial reaction time task methods***

The task was carried out using a bank of eight 9-hole chambers (Campden Instruments, UK), each enclosed in a sound attenuating chamber. Each of the equidistantly spaced holes across the array could present a visual stimulus and also register a nose-poke response by the breaking of an infrared beam. For this task, 4 of the 9 holes in the array (number 2, 4, 6 and 8) were blocked by masks. On the far side of the nose-poke array there was a food magazine, to which the reward was delivered by a peristaltic pump, via silicone tubing. Entrances to the food magazine were also recorded by means of an infrared. The chambers could be illuminated by lights in their side walls (“house lights), although all sessions were conducted in the dark apart from time out periods.

The mice were given a single 5-CSRTT session each day, where a session lasted for either a maximum of 60 trials or 20 minutes. Each trial at baseline conditions consisted of a stimulus presentation, pseudorandomly presented to one of the 5 locations, and a 5 s limited-hold period during which the stimulus was extinguished but the subject could still make a response and a 5 s inter-trial interval (ITI). A correct response (i.e. a nose-poke where the stimulus was illuminated) was rewarded by a 70 µl delivery of the reward, whereas following an incorrect response or an omission (i.e. no response) a 5 s time out period was initiated consisting of illumination of the house light. Responding during the ITI was defined as a “premature response” and initiated a time out and restarted the current trial.

The 5-CSRTT was commenced on the day following the final day of the reward preference test and broadly followed the protocol described by (1). In the first stage of shaping, the mice were habituated to the chambers for half an hour, during which reward was presented to the food magazine every 30 s. Mice completed a minimum of 3 sessions and then if they made >35 food magazine entries in two consecutive sessions they were moved to the next stage of training. Mice remained at this stage until they met the same criteria.

In the next stage of training, the mice performed a simplified version of the 5-CSRTT whereby only the central of the five holes was illuminated, with a stimulus duration of 60 s (1-CSRT task). Once performance reached criteria (>30 completed trials, >75% accuracy, <25% trial omissions) for two consecutive days, mice were moved on to the 5-CSRTT proper, with an initial stimulus duration of 60s. When mice met the above criteria, the stimulus duration was then decreased in the following way: 32 s, 16 s, 8 s, 4 s, 2 s, 1,8 s, 1.6 s, 1.4 s, 1.2 s, 1 s and 0.8 s. A stimulus duration of 0.8 s was considered baseline, and once reached, mice were kept on it for a minimum of five sessions and had to meet performance criteria for a minimum of two consecutive sessions, thus showing “stable” baseline performance before task manipulations were implemented to further study different aspects of attention and impulsivity. The average of the two final consecutive sessions before the first manipulation session was used to analyse performance at baseline conditions.

***RT-qPCR***

In preparation for the RT-qPCR the 1 μg of each sample’s RNA was reverse transcribed into cDNA with the EcoDry Premix double primed kit (Clontech), following the provided protocol. cDNA was diluted 1:10 in nuclease-free water and stored at -20 °C until further use. Primers for *Snord116, Snord115, Necdin, Mkrn3, Magel2* and *Ube3a* were used as targets of investigation. *Hprt, Gadph* and *B2m* were used for housekeeping genes with each sample as a positive control (see Supplementary Table 1). A reaction of 25 μL was prepared by adding 1.75 μL of each primer, 12.5 μL of 2X SensiMixSYBR No-ROX (Bioline), 5 μL of the sample and 4 μL of nuclease free water. Nuclease free water was added instead of a sample in each PCR run as a negative control. All samples and controls were ran in triplicates. The PCR was performed on a Corbet Rotor 30 Gene 6000 Real-Time PCR machine. The cycling conditions used for each reaction were: 1. 95°C 10 minutes, 2. 95°C 20 secs, 3. 60°C 20 secs, 4. 72°C 20 secs, repeat cycles 2-4 40 times, 5. 1°C increment increase from 50 to 99°C in order to generate melt curves, which were subsequently inspected to check that each primer pair was generating only one product.

The resulting data were averaged across the triplicates for each sample. ΔCt values were calculated by normalising expression data to the positive control -- subtracting the geometric mean of the housekeeping genes from the Ct value for each studied target gene in each sample. The ΔCt values for each condition were used for statistical analysis.

| **Gene** | **Primer sequence** | **Concentration** |
| --- | --- | --- |
| *Snord116* | ATCTAATGATGATTCCCAGTCAAACAT | 300 nM |
|  | TCACTCATTTTGTTCAGCTTTTCC | 300 nM |
| *Snord115* | ACAACCCACTGTCATGAAGAAAGG | 50 nM |
|  | CCTCAGCGTAATCCTATTGAGCAT | 900 nM |
| *Necdin* | ATGGTGCAGAAGCATCCTCAG | 300 nM |
|  | ATGGTGTGGAGATTGGTCAGC | 300 nM |
| *Mkrn3* | CCAATCAGTTGCTTAAGAAGTTGC | 300 nM |
|  | AAGAGCCAACGGTCATCAGAG | 300 nM |
| *Magel2* | GCATAGCAAGCCAGCCTCAG | 300 nM |
|  | GTAGACGAGCCTGTGGAGCCT | 50 nM |
| *Ube3a* | CAGACGTGACCATATTATAGATGATGC | 700 nM |
|  | CCACATACAACTGCTTCTTCAAGTCT | 700 nM |
| *Hprt* | GCGATGATGAACCAGGTTATGA | 300 nM |
|  | GCCTCCCATCTCCTTCATGA | 300 nM |
| *Gadph* | GAACATCATCCCTGCATCCA | 300 nM |
|  | CCAGTGAGCTTCCCGTTCA | 300 nM |
| *B2m* | TTCTGGTGCTTGTCTCACTGA | 300 nM |
|  | CAGTATGTTCGGCTTCCCATTC | 300 nM |

**Supplementary Table 1** RT-qPCR primer sequences for target and housekeeping genes

**SUPPLEMENTARY RESULTS**

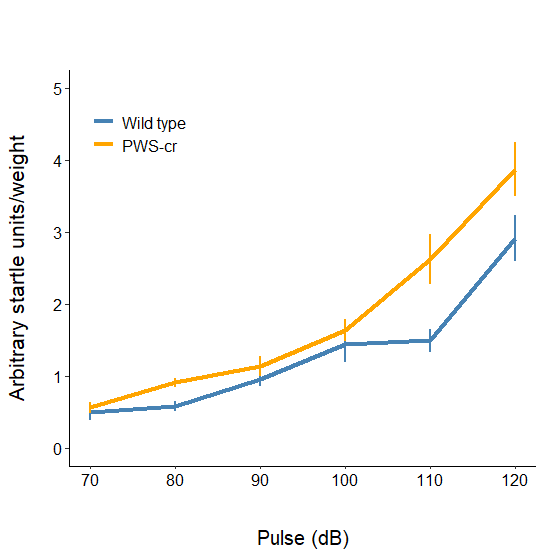

**Supplementary Figure 1.** Startle response to gradual increasing startle pulses (background noise 70dB) in PWS-cr and wild-type littermate mice. Data points are means for each group ±SEM.

ΔCt

|  | Wild type | PWS-IC |
| --- | --- | --- |
| Snord115 E13.5 | 0.137768 | 0.00369305 |
| Snord115 E18.5 | 1.67645 | 0.0649937 |
| Snord115 P0 | 7.26465 | 0.394545 |
| Snord116 E13.5 | 0.202273 | 0.00284798 |
| Snord116 E18.5 | 1.62048 | 0.0608103 |
| Snord116 P0 | 2.07974 | 0.152643 |
| Magel2 E13.5 | 0.0460154 | 0.000940225 |
| Magel2 E18.5 | 0.0231202 | 0.000356928 |
| Magel2 P0 | 0.0202934 | 0.00105502 |
| Mkrn3 E13.5 | 0.0803367 | 0.00198531 |
| Mkrn3 E18.5 | 0.0755087 | 0.000901803 |
| Mkrn3 P0 | 0.0914135 | 0.00365891 |
| Necdin E13.5 | 1.00997 | 0.0134981 |
| Necdin E18.5 | 1.51736 | 0.00834116 |
| Necdin P0 | 1.09022 | 0.0281895 |
| Ube3a E13.5 | 1.36443 | 1.65909 |
| Ube3a E18.5 | 1.01349 | 1.73114 |
| Ube3a P0 | 0.933107 | 2.32948 |

**Supplementary Figure 2.** Expression of the genes from the PWS locus in whole brain tissue at two developmental and one postnatal stage. RT-qPCR data validated the expression of the PWS-IC model, by showing the expected loss of expression of the PEGs *Snord115, Snord116* and *Necdin,* and an overexpression of the MEG *Ube3a*, particularly at 18.5E stage of foetal mouse development and at birth (bars indicate mean, error bars indicate SEM).

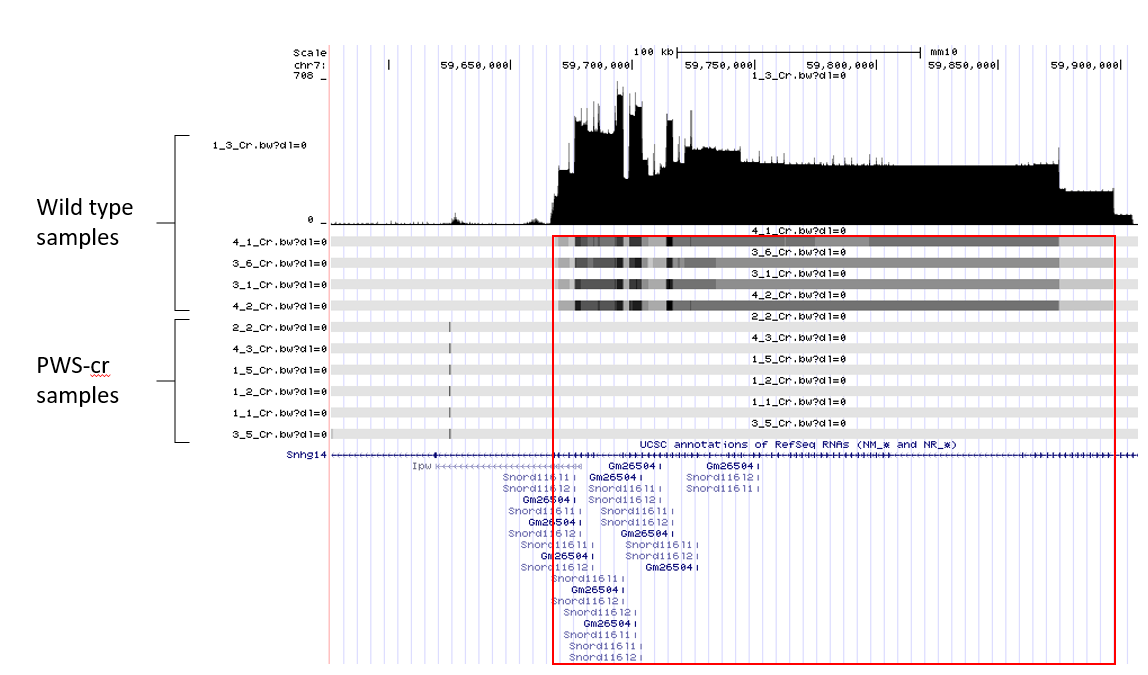

**Supplementary Figure 3.** UCSC browser track image of STAR generated transcript counts at the critical interval of PWS-cr mouse model and their wild type littermates shows the deletion in the PWS-cr mice spans all the copies of *Snord116,* as well as most of the exons of the *Ipw* non-coding RNA.
